# Tensor-decomposition–based unsupervised feature extraction in single-cell multiomics data analysis

**DOI:** 10.1101/2021.08.25.457731

**Authors:** Y-h. Taguchi, Turki Turki

## Abstract

Analysis of single-cell multiomics datasets is a novel topic and is considerably challenging because such datasets contain a large number of features with numerous missing values. In this study, we implemented a recently proposed tensor-decomposition (TD)–based unsupervised feature extraction (FE) technique to address this difficult problem. The technique can successfully integrate single-cell multiomics data composed of gene expression, DNA methylation, and accessibility. Although the last two have large dimensions, as many as ten million, containing only a few percentages of non-zero values, TD-based unsupervised FE can integrate three omics datasets without filling missing values. Together with UMAP, which is used frequently when embedding single-cell measurements into two-dimensional space, TD-based unsupervised FE can produce two-dimensional embedding coincident with classification when integrating single-cell omics datasets. Genes selected based on TD-based unsupervised FE were also significantly related to reasonable biological roles.

## 1. Introduction

Single-cell multiomics data analysis is challenging [1]. There are multiple reasons for this issue. First, it inevitably includes too many missing values. In the usual high throughput sequencing (HTS), the so called depth can compensate for this problem. Nevertheless, because of very limited amount of RNA retrieved from individual cells available, ‘depth’ cannot resolve this missing value problems. Second, too many missing values result in apparent diversity. The primary purpose of single-cell analysis is to identify the diversity of individual cells that cannot be recognized by the tissue-level HTS. Although missing values are random, apparently very variant profiles appear from a single profile, which can be recognized if there is a large enough number of reads available. This compels researchers to distinguish between true biological diversity and apparent diversity caused by missing values [2].

Finally, single-cell analysis is computationally challenging. Because there are not many samples in the standard HTS, even if the number of features is large, the overall required computational resources decided by the product between the number of features and the number of samples are very limited. Nonetheless, since the number of samples that is the same as that of cells can be huge in single-cell analysis, single-cell analysis can be computationally very challenging.

To resolve these difficulties, we employed tensor-decomposition (TD)–based unsupervised feature extraction (FE) [3]. Prior to applying TD to multiomics datasets, singular value decomposition (SVD) was applied to individual omics profiles such that individual omics profiles have common *L* singular value vectors. Then, *K* omics profiles are formatted as an *L* × *M* × *K* dimensional tensor, where *M* is the number of single cells. Then, higher-order singular value decomposition (HOSVD), which is a type of TD, is applied to the tensor. UMAP applied to singular value vectors attributed to single cells by HOSVD successfully generated two dimensional embedding, coincident with known classification of single cells.

## 2. Materials and Methods

### 2.1. Gene expression profiles

Two single-cell multiomics datasets were downloaded from GEO using the following two GEO IDs:

#### 2.1.1. GSE154762:data set 1

The multiomics dataset [4] retrieved from GEO ID GSE154762, which is denoted as dataset 1 in this study, is composed of 899 single cells for which gene expression, DNA methylation, and DNA accessibility were measured. These single cells represent human oocyte maturation (Table 1). For gene expression, the file “GSE154762_hO_scChaRM_count_matix was downloaded from the supplementary file of GEO and was loaded onto R [5] using `read.table` function in R. For DNA methylation and DNA accessibility, 899 files with the extensions “WCG.bw” and “HCG.bw” were downloaded from supplementary files of GEO and were loaded onto R using the `import` function in rtracklayer [6] package in R.

**Table 1.** The number of single cells within individual cell types included in data set 1.

| FGO | GO1 | GO2 | Granulosa | Immune | MI | MII | StromaC1 | StromaC2 |
| --- | --- | --- | --- | --- | --- | --- | --- | --- |
| 81 | 40 | 46 | 93 | 20 | 155 | 90 | 189 | 185 |

#### 2.1.2. GSE121708:data set 2

The multiomics dataset [7] retrieved from GEO ID GSE154762, which is denoted as dataset 2 in this study, is composed of 852 single cells for which DNA methylation and DNA accessibility were measured as well as 758 single cells for which gene expression was measured. These single cells represent the four time points of the mouse embryo (Table 2). For gene expression, the file “GSE121650_rna_counts.tsv.gz” was downloaded from the supplementary file of GEO and was loaded onto R using the `read.table` function in R. For DNA methylation and DNA accessibility, 852 files with the extensions “met.tsv.gz” and “acc.tsv.gz” were downloaded from supplementary file of GEO and were loaded onto R using the `read.table` function in R.

**Table 2.** The number of single cells at four embryonic time points included in dataset 2. For E7.5, the gene expression profiles of 296 single cells were measured.

| E4.5-5.5 | E6.5 | E6.75 | E7.5 |
| --- | --- | --- | --- |
| 267 | 98 | 97 | 390(296) |

### 2.2. Pre-processing of DNA methylation profiles

First, we collected genomic positions for which at least one measurement was performed for at least one single cell (i.e., union). Then, for each genomic position, three integers, −1, 0, and 1, were assigned. When the genomic position was measured in a single-cell and its state was methylated (non-methylated), we attributed 1 (−1) to the genomic position of the single cell. Otherwise (i.e., missing observation), we attributed 0 to the genomic transition in a single cell. 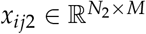 is stored as a sparse matrix object using the Matrix [8] package in R because of the large *N*_2_.

### 2.3. Pre-processing of DNA accessibility

First, we divided the whole genome into 200 nucleotide regions, and DNA accessibility was summed up within individual regions. These values, which show the summation of DNA accessibility within individual regions, are regarded as DNA accessibility at the individual 200 nucleotide regions, each of which is supposed to approximately correspond to a single nucleosome that is composed of 140 length DNA that wraps around histones and 80 length of linker DNA. In this study, these 200 nucleotide regions are called as “nucelosome regions”. 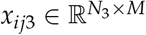 was stored as a sparse matrix object using the Matrix package in R because of the large *N*_3_.

### 2.4. TD-based unsupervised FE

#### 2.4.1. Reduction of feature dimensions

Here, feature denotes gene expression, DNA methylation, or DNA accessiblity. Because the features of these three datasets differ from one another, we first applied SVD to these features. Suppose 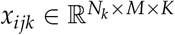 represents the value of the *i*th feature (expression of the *i*th gene, methylation of the *i*th genomic location, or DNA accessibility of the *i*th nucleosome region) at the *k*th single cell of the *k*th omics datae (1 ≤ *k* ≤ *K* = 3, *k* = 1:gene expression, *k* = 2:DNA methylation, and *k* = 3:DNA accessibility). Applying SVD to *x_ijk_*, and we get

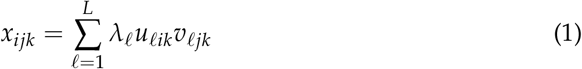

where λ_*ℓ*_ is the *ℓ*th singular value, and *u_ℓ_i_k_* and *υ_ℓjk_* are the *i*th and *j*th components of the *ℓ*th left and right singular value vectors, respectively. Then, *x_ijk_* is transformed to 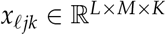 to have the same (common) feature dimension, *L*, independent of *k*, as

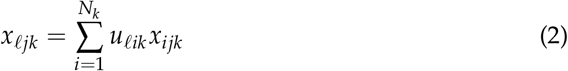

#### 2.4.2. Data normalization

Prior to applying SVD to the individual omics profiles in these two datasets, *x_ijk_*, (*k* = 2, 3), that is, DNA methylation and accessibility, of dataset 1 is normalized such that

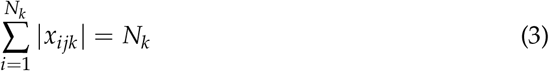

whereas *x_ij1_*, i.e., gene expression, is normalized such that

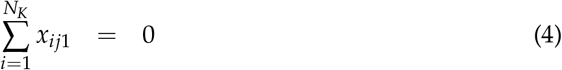

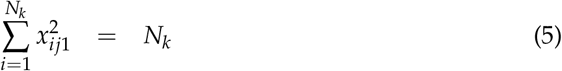

for datasets 1 and 2. The reason why DNA methylation and accessibility of dataset 2 are not normalized is because ∑_*i*_|*x_ijk_*|, (*k* = 2, 3) is very small in some single cells in dataset 2. Thus, applying normalization adds significant weight to these single cells with fewer observations and drastically skewed outcomes. To avoid this problem, *x_ijk_*, (*k* = 2, 3) of dataset 2 was not normalized.

#### 2.4.3. TD applied to dimension reduced multiomics data sets

HOSVD [3] was applied to the tensor, *x_ℓjk_*, and we got

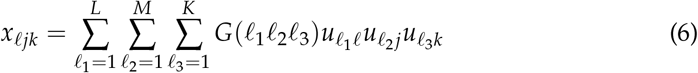

where 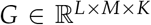 is the core tensor that represents the contribution of 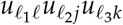 to *x_ℓjk_*. 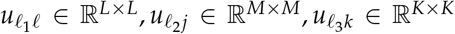 are singular value matrices and are orthogonal matrices.

### 2.5. Categorical regression

For categorical regression to test the coincidence between classification shown in Tables 1 or 2 and singular value vectors attributed to the *j*th single cells, we performed categorical regression

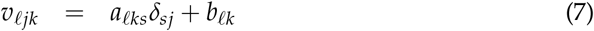

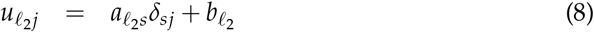

where *s* denotes one of the classifications shown in Table 1 or 2, *a_ℓks_, b_ℓk_, a_ℓ_2_ks_, b_ℓ_2_k_* are regression coefficients, and *δ_js_* takes the value of 1 when the *j*th single cell belongs to the sth classification and 0 otherwise. Categorical regression was performed using the ls function in R. The obtained *P*-values were corrected using the Benjamini-Hochberg (BH) criterion [3]. *ℓ*s or *ℓ*_2_s associated with adjusted *P*-values less than 0.01 are regarded to be coincident with classification.

### 2.6. UMAP

Two-dimensional embedding was performed by UMAP [9]. The `umap` function implemented in R was used.

### 2.7. Gene selection

After identifying which *u_ℓ_2_j_* coincides with the classification, we need to identify which *u_ℓ_1_ℓ_* is associated with the selected *u_ℓ_2_j_* by investigating |*G*(*ℓ*_1_*ℓ*_2_*ℓ*_3_)|; *ℓ*_1_s with a larger |*G*| with the selected *ℓ*_2_ are regarded to be coincident with the classification. Then, the selected *u_ℓ_1_ℓ_* is converted back to *u_ℓ_1_i_1__* attributed to genes as

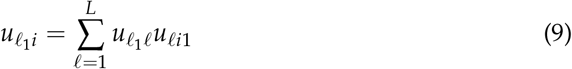

*P*-values can be attributed to genes, *i*, assuming *u_ℓ_1_i_* obeys a multiple Gaussian distribution (null hypothesis) as

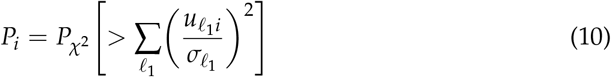

where the summation is taken over only the selected *ℓ*_1_s, and *P*_*χ*^2^_ [> *x*] is the cumulative *χ*^2^ distribution, where the argument is larger than *x*, and *σ_ℓ_1__* is the standard deviation. *P_i_*s are corrected by the BH criterion [3], and *i*s associated with adjusted *P_i_* less than 0.01 are selected.

## 3. Results

### 3.1. Data set 1

We got 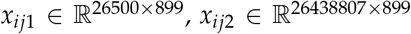, and 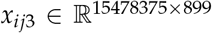. SVD was applied to *x_ijk_* with *L* = 10, as in Eq. (1). For *x_ijk_, k* = 2, 3, SVD was performed using the `irlba` function in the irlba package [10] in R because of the large *N_k_, k* = 2, 3 as many as 10 million. Then, HOSVD was applied to *x_ℓjk_*, as in eq. (2).

One possible validation to check whether the above procedure works properly is to check whether *υ_ℓjk_* and *u_ℓ_2_j_* are coincident with the classification shown in Table 1. Because the above procedure is fully unsupervised, it is unlikely that *υ_ℓjk_* and *υ_ℓ_2_j_* are accidentally coincident with the classification. To quantitatively validate the coincidence between the classification and *υ_ℓjk_* or *u_ℓ_2_j_*, we applied categorical regression (see §2.5).

Table 3 shows the number of singular value vectors coincident with the classification shown in Table 1. When SVD was applied to individual omics data, because *L* = 10, the number of singular value vectors was 10 as well. When *K* omics datasets are integrated and HOSVD is applied, the number of singular value vectors is *KL* = 10*K*. Thus, when DNA methylation and accessibility are integrated, 20 singular value vectors are available. When all three omics data are integrated, 30 singular value vectors are available. It is obvious that for all five cases, at least one singular value vector is coincident with the classification. Thus, our strategy is essentially successful.

To further validate the successful integration of singular value vectors, we applied UMAP to 20 or 30 singular value vectors obtained by HOSVD (Fig. 1).

**Figure 1.**
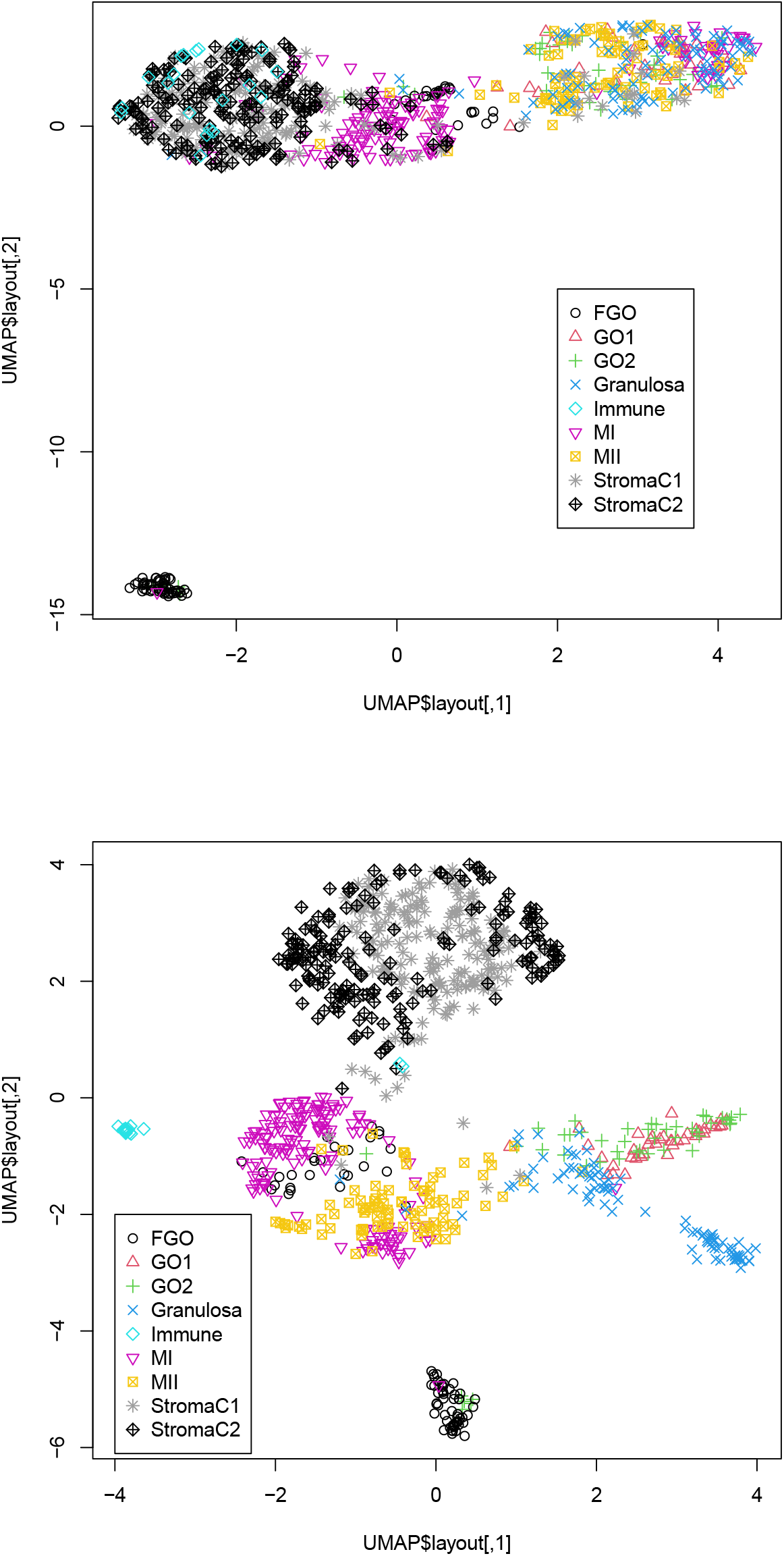
Two-dimensional embedding of singular value vectors, *u_ℓ_2_j_*, computed by HOSVD applied to *x_ℓjk_* in dataset 1 (Table 3). Upper: *u_ℓ_2_j_*, 1 ≤ *ℓ*_2_ ≤ 20 when only DNA methylation and accessibility (*k* = 2, 3) are integrated. Lower: *u_ℓ_2_j_*, 1 ≤ *ℓ*_2_ ≤ 30 when all three omics data points (1 ≤ *k* ≤ 3) are integrated. Default settings other than custom.config$n_neighbors=100 were used.

**Table 3.** Number of singular value vectors coincident with classification shown in Table 1.

| Adjusted<br>*P*-value | gene<br>expression | SVD ($v_{\ell jk}$)<br>DNA<br>methylation | DNA<br>accessibility | HOSVD ($u_{\ell_2 j}$)<br>DNA methylation<br>and accessibility | All |
| --- | --- | --- | --- | --- | --- |
| < 0.01 | 10 | 7 | 1 | 10 | 18 |
| $\geq 0.01$ | 0 | 3 | 9 | 10 | 12 |

It is obvious that the integration of all three omics datasets (lower) is more coincident with classification than that of integration of two omics datasets, DNA methylation, and accessibility (upper). This suggests the usefulness of integrating the three omics datasets. In fact, single omics data cannot provide two-dimensional embedding coinciding with classification (Fig. S1).

We also attempted to validate biological outcomes when all three omics datasets were integrated. We selected 47 genes associated with adjusted *P_i_* less than 0.01, as described in §2.7 using *u_1i_* because *u_1i_* is associated with the largest

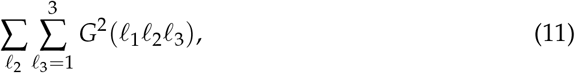

where the summation of *ℓ*_2_ is taken over only 18 *ℓ*_2_s coincident with the classification (Table 3). The selected 47 genes (Data S1) were uploaded to Enrichr [11].

Forty-seven genes were enriched by H3K36me3 based on “ENCODE Histone Modifications 2015”; H3K36m3 is known to play critical roles during oocyte maturation [12]. Forty-seven genes were also targeted by MYC based on “ENCODE and ChEA Consensus TFs from ChIP-X”; Myc is known to play critical roles in oogenesis [13]. Forty-seven genes were also targeted by TAF7 based on “ENCODE and ChEA Consensus TFs from ChIP-X” and “ENCODE TF ChIP-seq 2015”; TAF7 is known to play critical roles during oocyte growth [14]. Forty-seven genes are also targeted by ATF2 based upon “ENCODE and ChEA Consensus TFs from ChIP-X”; expression of ATF2 is known to be altered during oocyte dvelopment [15]. This suggests that our strategy correctly captures regulation-related parts (full data of enrichment analysis can be available as Data S1).

### 3.2. Data set 2

To confirm that the success in the previous section is not accidental, we applied the same procedure to dataset 2 as well. We got 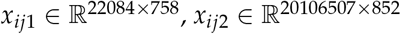, and 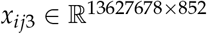. SVD was applied to *x_ijk_* with *L* = 10, as in Eq. (1). For *x_ijk_, k* = 2, 3, SVD was performed using the irlba function in the irlba package [10] in R because of the large *N_k_, k* = 2, 3 of as many as ten millions. Then, HOSVD was applied to *x_ℓjk_*, as in eq. (2). Because *N*_1_ = 758 < *N*_2_ = *N*_2_ = 852, when HOSVD is applied to *x_ℓjk_* composed of all three omics datasets, only 758 single cells shared with all *x_ijk_* are considered. As described in the previous section, we first validated the coincidence between singular value vectors attributed to single cells, that is, *υ_ℓj_* and *u_ℓ_2_ℓk_*, and classification in Table 2 (Table 4).

**Table 4.**
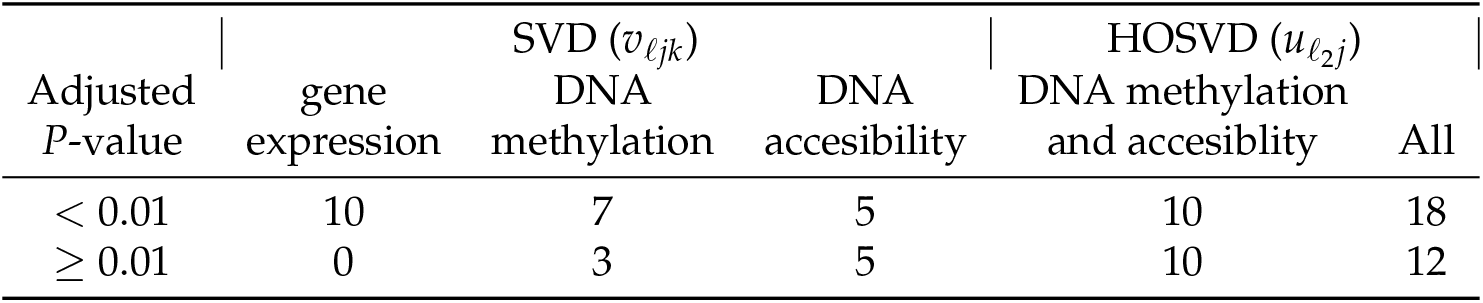
Number of singular value vectors coincident with classification shown in Table 2.

| Adjusted<br>$P$-value | gene<br>expression | SVD ($v_{\ell jk}$) | | HOSVD ($u_{\ell_2 j}$) | |
| --- | --- | --- | --- | --- | --- |
|  |  | DNA<br>methylation | DNA<br>accessibility | DNA methylation<br>and accessibility | All |
| $< 0.01$ | 10 | 7 | 5 | 10 | 18 |
| $\geq 0.01$ | 0 | 3 | 5 | 10 | 12 |

The coincidence between the singular value vectors and the classification in Table 4 is even better than those in Table 3. Thus, it is unlikely that the superior outcome in Table 3 is purely accidental. To further validate the successful integration of singular value vectors, we applied UMAP to 20 or 30 singular value vectors obtained by HOSVD (Fig. 2).

**Figure 2.**
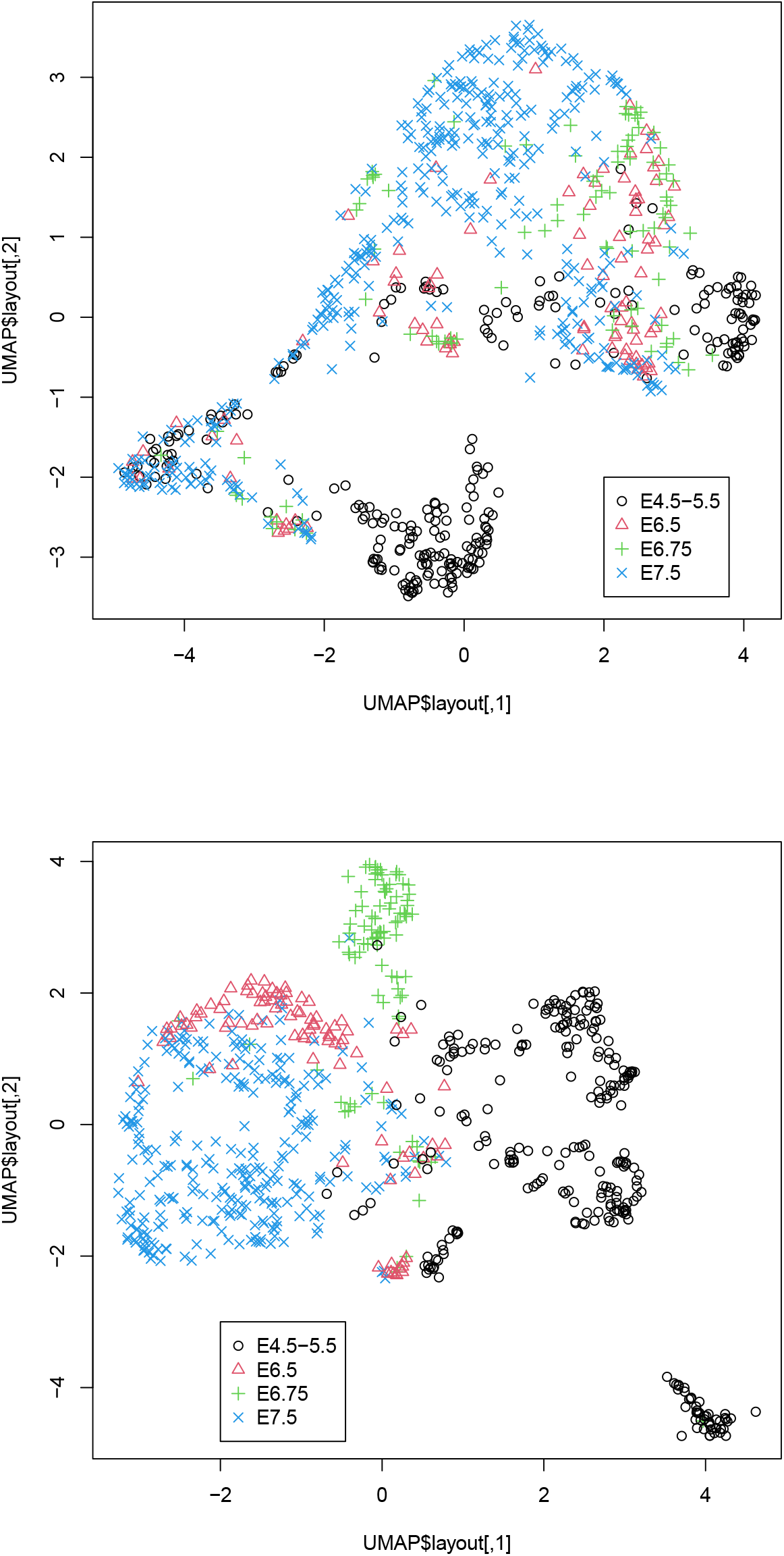
Two-dimensional embedding of singular value vectors, *u_ℓ_2_j_*, computed by HOSVD applied to *x_ℓjk_* in dataset 2 (Table 4). Upper: *u_ℓ_2_j_*, 1 ≤ *ℓ*_2_ ≤ 20 when only DNA methylation and accessibility (*k* = 2, 3) are integrated. Lower: *u_ℓ_2_j_*, 1 ≤ *ℓ*_2_ ≤ 30 when all three omics data points (1 ≤ *k* ≤ 3) are integrated. Default settings other than custom.config$n_neighbors=100 were used.

It is obvious that the integration of all three omics datasets (lower) is more coincident with classification than that of integration of two omics datasets, DNA methylation, and DNA accessibility (upper), as can be seen in Fig. 2. This again confirms the usefulness of integrating the three omics datasets. In fact, single omics data cannot provide two-dimensional embedding coinciding with classification (Fig. S2).

We also attempted to validate biological outcomes when all three omics datasets were integrated. We selected 175 genes associated with adjusted *P_i_* less than 0.01, as described in §2.7 using *u_1_i* because *u_1_i* is associated with the largest

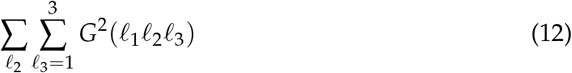

where the summation of *ℓ*_2_ is taken over only 18 *ℓ*_2_s coincident with the classification (Table 4). The selected 175 genes (Data S2) were converted to gene symbols by DAVID [16] gene ID converter and were uploaded to Enrichr.

One hundred and seventy-five genes were enriched by H3K36me3 based on “ENCODE Histone Modifications 2015”; H3K36m3 is known to play critical roles during gastrulation [17]. One hundred and seventy-five genes were also targeted by MYC based on “ENCODE and ChEA Consensus TFs from ChIP-X”; Myc is also known to play critical roles in gastrulation [18]. One hundred and seventy-five genes were also targeted by TAF7 based on “ENCODE and ChEA Consensus TFs from ChIP-X” and “ENCODE TF ChIP-seq 2015”; TAF7 is known to play critical roles during gastrulation [19]. One hundred and seventy-five genes were also targeted by ATF2 based on “ENCODE and ChEA Consensus TFs from ChIP-X”; expression of ATF2 is known to be maintained during gastrulation [20]. This suggests that our strategy correctly captures regulation-related parts (full data of enrichment analysis is available as Data S2).

These two examples, application to datasets 1 and 2, demonstrate the usefulness of the present strategy to integrate single-cell multimics datasets composed of gene expression, DNA methylation, and accessibility.

## 4. Discussion

In this study, we demonstrated the usefulness of our strategy when it is applied to the integrated analysis of single-cell multiomics datasets composed of gene expression, DNA methylation, and DNA accessibility. One might wonder if other more popular methods can achieve similar performance because our strategy is useless if others can perform comparatively. There are several advantages of our method, which other methods do not have.

First, we do not have to fill the missing values with non-zero values. Single-cell measurements are usually associated with a large number of missing values (Table 5).

**Table 5.**
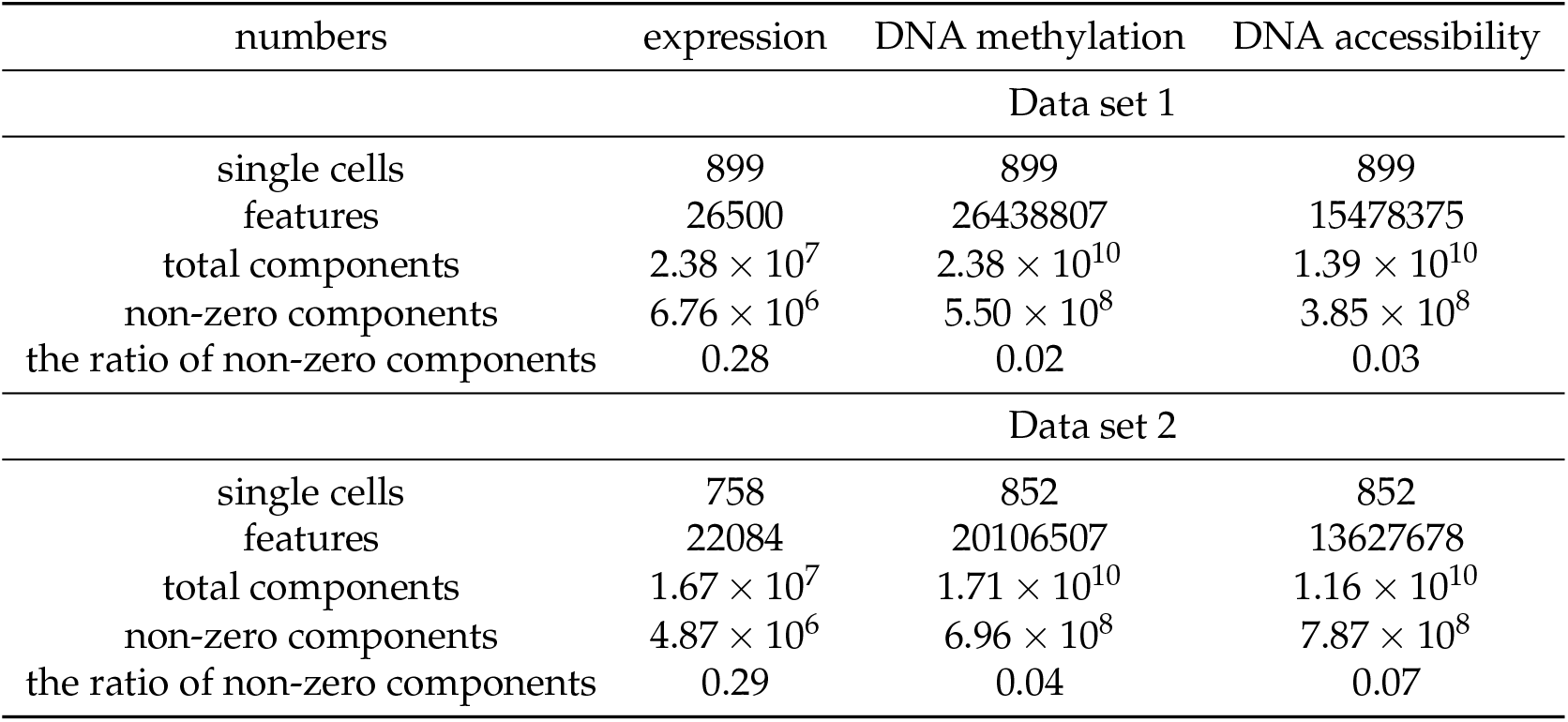
Number of single cells, features, non-zero components, and its ratios.

| numbers | expression | DNA methylation | DNA accessibility |
| --- | --- | --- | --- |
| Data set 1 |  |  |  |
| single cells | 899 | 899 | 899 |
| features | 26500 | 26438807 | 15478375 |
| total components | $2.38 \times 10^7$ | $2.38 \times 10^{10}$ | $1.39 \times 10^{10}$ |
| non-zero components | $6.76 \times 10^6$ | $5.50 \times 10^8$ | $3.85 \times 10^8$ |
| the ratio of non-zero components | 0.28 | 0.02 | 0.03 |
| Data set 2 |  |  |  |
| single cells | 758 | 852 | 852 |
| features | 22084 | 20106507 | 13627678 |
| total components | $1.67 \times 10^7$ | $1.71 \times 10^{10}$ | $1.16 \times 10^{10}$ |
| non-zero components | $4.87 \times 10^6$ | $6.96 \times 10^8$ | $7.87 \times 10^8$ |
| the ratio of non-zero components | 0.29 | 0.04 | 0.07 |

Although gene expression profiles are associated with a relatively small number of missing components, more than 70 % were missing. For DNA methylation and accessibility, the situation is very difficult to treat. Only a few percentages of components have values, while the rest are missing values. To address this problem, especially for DNA methylation and accessibility, heavy pre-processing is usually required. For example, for dataset 1, statistical tests are applied and regions associated with significant *P*-values are selected [4], which reduces the number of features attributed to DNA methylation and accessibility. Because such a statistical test automatically filters out regions filled with missing values, the ratio of non-zero components is also reduced as a result. For dataset 2, the authors restricted the features to only the most variable ones (typically, ~ 10^3^) and occasionally filled missing components with Bayesian models [7]. These procedures inevitably introduce arbitrariness of outcomes, as pre-processing data might affect the outcome. In contrast to these arbitrary procedures, our method is almost unsupervised. We did not select any features or fill in the missing values. Despite these fully unsupervised strategies, our results are highly coincident with the classification (Tables 3 and 4 and Figs. 1 and 2). From this perspective, our strategy is superior to the other methods.

Second, our method can deal with massive datasets. For example, although integrated analysis of multiomics data was performed using multiomics factor analysis (MOFA) [21] in the original studies [4,7] of datasets 1 and 2, MOFA cannot accept *x_ijk_* in this study as inputs because MOFA does not implement sparse matrix architecture. During the computation of MOFA, zero values must be filled with non-zero values to evaluate the convergence; this results in a dense matrix that cannot be stored in the computer memory because the number of components of DNA methylation and accessibility are too large to store them as they are (Table 5). In our computation, we can apply SVD to these large datasets while keeping them in a sparse matrix format using the irlba package implemented in R. SVD not only reduces the number of features to *L* but also fills missing values. Thus, we can manage a large matrix as it is in our implementation.

Third, our method is free from dividing weight between multiomics data sets; how to weigh individual omics data must be decided based on some criteria outside the datasets available. Nevertheless, in our implementation, the weight of individual omics data is represented by *u_ℓ_3_k_*, which is automatically decided by simply applying HOSVD to a multiomics dataset. Thus, from this perspective, our strategy is outstanding.

Although we have shown that the integration of all three omics data are superior to that of integration of DNA methylation and accessibility (Figs. 1 and 2), one might wonder if integration of gene expression and DNA methylation or DNA accessibility might be comparative with that of all thee omics data sets. In order to deny this possibility, we also considered these combinations of two of three omics data sets (Figs. S3 and S4). Although the integration of gene expression and DNA accessibility in dataset 1 (Fig. S3(B)) is comparative with that of all three omics data, neither integration of gene expression and DNA memthylation (Fig. S4(A)) nor that of gene expression and DNA accessibility (Fig. S4(B)) is comparative with that of all three omics data in dataset 2. Thus, it is obvious that only integaration of three omics datasets can give us UMAP embeddinng coincident with classification regardless to the dataset considerd.

As for the comparisons with other methods, as mentioned above, no methods implemented with sparse matrix architecture and applicable to multiomics datasets exist to our knowledge. Thus, we could not compare our performance with other methods.

Prospective uses of our methods are as follows. First of all, it can integrate gene expression profiles, DNA methylation and accessibility in single cell measurements with applying no pre-processing; this enables researchers to obtain the reasonable results without struggling on how to convert raw data into treatable formats. In addition to this, since it can save memories required for analysing single cell multiomics data sets, more researchers who cannot have massive computational facilities can analyze massive single cell measurements.

## 5. Conclusions

In this study, we propose a method for applying TD-based unsupervised FE to single-cell multiomics datasets composed of gene expression, DNA methylation, and DNA accessibility. Together with UMAP, the proposed method successfully integrated a multiomics dataset and generated two-dimensional embedding of single cells coincident with the classification. The implementation requires neither filling missing values nor massive CPU memory to store multiomics datasets of single cells and can deal with DNA methylation and accessibility with ten million features. The present implementation is very promising and can be a *de facto* standard method to integrate single-cell multiomics datasets composed of gene expression, DNA methylation, and DNA accessibility.

## Author Contributions

YHT planned the research, performed analyses. YHT and TT have evaluated the results, discussions, outcomes and wrote and reviewed the manuscript.

## Data Availability Statement

Data used in this study is available in GEO ID GSE154762 and GSE121708. Sample R source can be found at https://github.com/tagtag/scMultiR

## Abbreviations

The following abbreviations are used in this manuscript:

BH: Benjamini-Hochberg
FE: feature extraction
HOSVD: higher order singular value decomposition
HTS: high throughput sequencing
MOFA: Multi-Omics Factor208Analysis
SVD: singular value decomposition
TD: tensor decomposition

