## Supplementary material for "Tensor-decomposition–based unsupervised feature extraction in single-cell multiomics data analysis": Data S1, Data S2 and Figs. S1 to S4: Supplementary_Figures.pdf

(A)

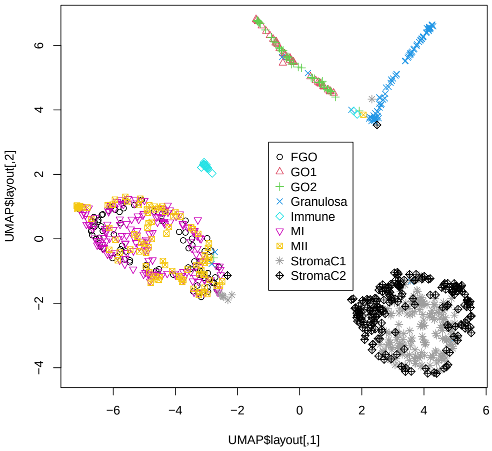


(B)

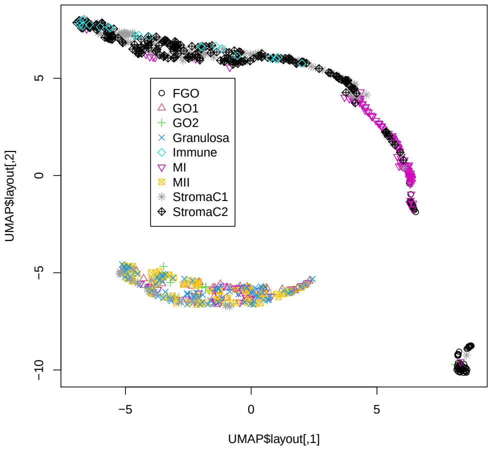


(C)

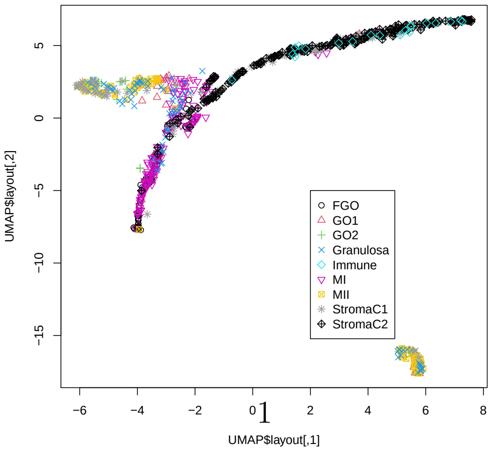




Figure S1: UMAP embedding of single omics data for data set 1. (A) gene expression (B) DNA methylation (C) DNA accessibility. Default setting other than custom.config$n_neighbors=100 are used.

(A)

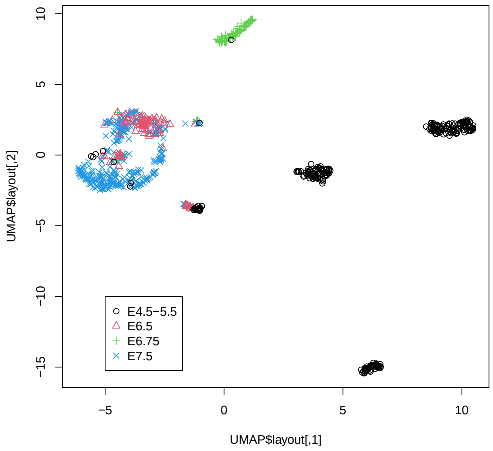


(B)

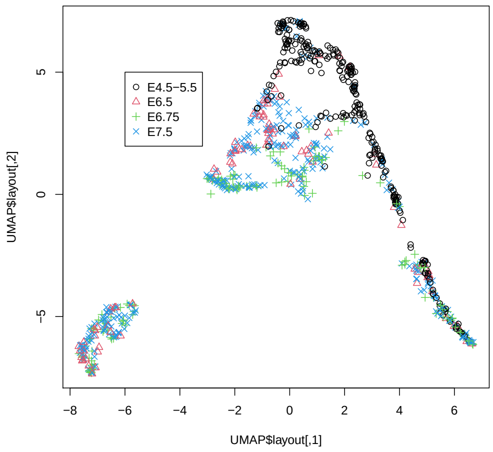


(C)

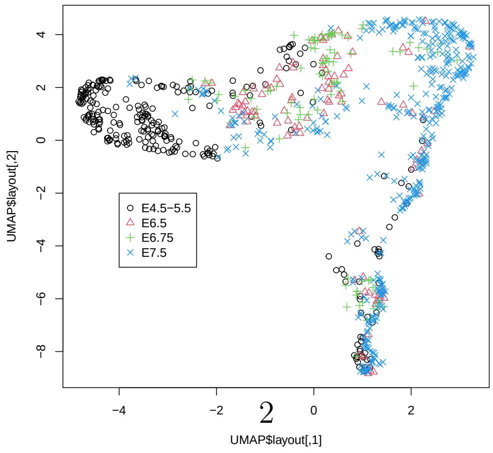




Figure S2: UMAP embedding of single omics data for data set 2. (A) gene expression (B) DNA methylation (C) DNA accessibility. Default setting other than custom.config$n_neighbors=100 are used.

(A)

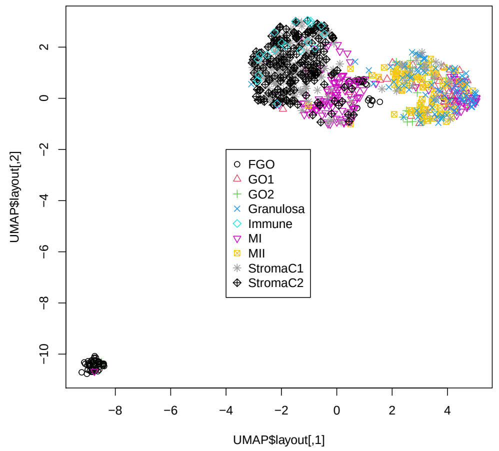


(B)

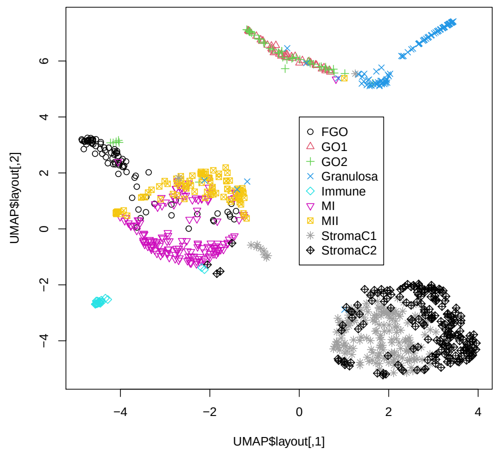


Figure S3: Two-dimensional embedding of singular value vectors, $u_{\ell_2 j}$, computed by HOSVD applied to $x_{\ell jk}$ in dataset 1. (A) $u_{\ell_2 j}, 1 \leq \ell_2 \leq 20$ when only gene expression and DNA methylation ($k = 1, 2$) are integrated. (B) $u_{\ell_2 j}, 1 \leq \ell_2 \leq 20$ when only gene expression and DNA accessibility ($k = 1, 3$) are integrated. Default settings other than custom.config$n_neighbors=100 were used.

(A)

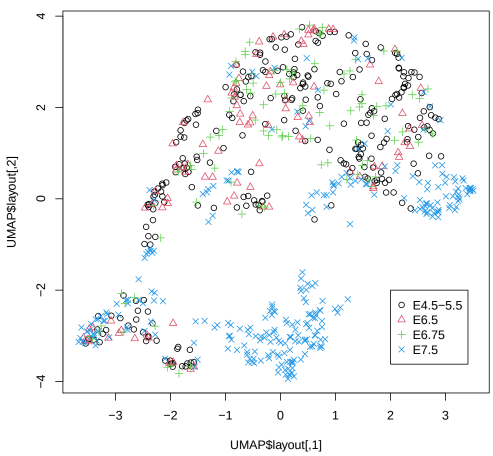


(B)

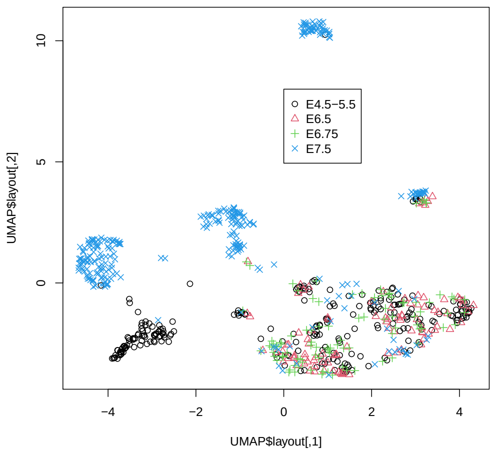


Figure S4: Two-dimensional embedding of singular value vectors, $u_{\ell_2 j}$, computed by HOSVD applied to $x_{\ell jk}$ in dataset 2. (A) $u_{\ell_2 j}, 1 \leq \ell_2 \leq 20$ when only gene expression and DNA methylation ($k = 1, 2$) are integrated. (B) $u_{\ell_2 j}, 1 \leq \ell_2 \leq 20$ when only gene expression and DNA accessibility ($k = 1, 3$) are integrated. Default settings other than custom.config$n_neighbors=100 were used.
